# Lysosome damage triggers direct ATG8 conjugation and ATG2 engagement via CASM

**DOI:** 10.1101/2023.03.22.533754

**Authors:** Jake Cross, Joanne Durgan, David G McEwan, Oliver Florey

## Abstract

Cells harness multiple pathways to maintain lysosome integrity, a central homeostatic process. Damaged lysosomes can be repaired, or targeted for degradation by lysophagy, a selective autophagy process involving ATG8/LC3. Here, we describe a parallel ATG8/LC3 response to lysosome damage, mechanistically distinct from lysophagy. Using a comprehensive series of biochemical, pharmacological and genetic approaches, we show that lysosome damage induces rapid <u>C</u>onjugation of <u>A</u>TG8s to <u>S</u>ingle <u>M</u>embranes (CASM). ATG8 proteins are recruited directly onto damaged membranes, independently of ATG13/WIPI2, and conjugated to PS, as well as PE, a molecular hallmark of CASM. Lysosome damage drives V-ATPase V0-V1 association, and direct recruitment of ATG16L1, dependent on K490 (WD40-domain), and sensitive to *Salmonella* SopF. Lysosome damage-induced CASM is associated with the formation of dynamic LC3A-positive tubules, and promotes robust LC3A engagement with ATG2, a lipid transfer protein central to lysosome repair. Together, our data identify direct ATG8 conjugation as a rapid response to lysosome damage, with important links to lipid transfer and dynamics.

## Introduction

The lysosome is an acidic, catabolic organelle, which plays a central role in the degradation of extra- and intra-cellular cargoes (Ballabio and Bonifacino, 2020) to maintain cellular homeostasis and health. Lysosomes also play a vital signalling function, acting as hubs to sense, integrate and respond to changes in nutrient status, metabolic signals and cellular stress (Perera and Zoncu, 2016). Given these fundamental functions, maintenance of lysosomal integrity is of paramount importance for cellular and organismal homeostasis. Many agents and conditions can threaten this integrity by damaging late endosomal or lysosomal compartments, through partial lysosomal membrane permeabilization (LMP), or full rupture of lysosomes. Damage can be caused by endocytosed materials, such as silica or urate crystals, that mechanically rupture the membrane, as well as neurotoxic aggregates, pathogens and membrane lipid changes associated with aging (Wang et al., 2018; Zhen et al., 2021). To defend against such perturbations, cells have developed sophisticated defence mechanisms, that together comprise the Lysosomal Damage Response (LDR).

Small-scale lysosome damage can be repaired via an ESCRT-mediated membrane remodelling process, that is dependent on Ca2+ release (Radulovic et al., 2018; Scheffer et al., 2014; Skowyra et al., 2018). Additionally, a parallel repair pathway, termed phosphoinositide-initiated membrane tethering and lipid transport (PITT), has recently been described (Tan and Finkel, 2022). During PITT, PI4K2A is recruited to damaged lysosomes, to increase the level of PI4P. This in turn engages ORP-VAPA lipid transfer complexes, to form ER contact sites which transfer PS and cholesterol to the lysosome membrane (Radulovic et al., 2022). Subsequently, the lipid transfer protein, ATG2, is recruited, supplying further lipids to repair the damaged lysosome. Another protein, LRRK2, is also recruited to stressed lysosomes, where it plays an important role in restoring integrity (Eguchi et al., 2018; Herbst et al., 2020), through the generation of lysosome tubules, in a process termed LYsosomal Tubulation/sorting driven by LRRK2 (LYTL) (Bonet-Ponce et al., 2020). These repair mechanisms act in parallel to ensure robust protection of the lysosomal system.

In the case of more profound membrane damage, lysosomes can instead be cleared by a selective autophagy process called lysophagy (Maejima et al., 2013). Lysophagy involves the recognition of luminal galectins, which when exposed to the cytosol promote the recruitment of the ULK/ATG13/FIP200 autophagy initiation complex. This complex acts in concert with Vps34 and PI3P generation to promote the *de novo* formation of a double-membrane autophagosome that sequesters the damaged lysosome (Maejima et al., 2013). A key feature of this autophagosome formation is the involvement of a ubiquitin-like conjugation system, including ATG16L1, to target the conjugation of ATG8 proteins (LC3A/B/C and GABARAP/L1/L2 families), to phosphatidylethanolamine (PE) in the forming autophagosome membrane (Fujita et al., 2013; Ichimura et al., 2000). ATG8 proteins act with WIPI4 to recruit ATG2, initiating contacts with the ER and transferring lipids to support autophagosome expansion (Bozic et al., 2020; Osawa et al., 2019; Valverde et al., 2019). ATG2 thus functions in both the PITT pathway and during classical lysophagy, as a central player in lysosome homeostasis.

A subset of autophagy proteins can also function in a parallel “non-canonical autophagy” pathway, defined by the <u>C</u>onjugation of <u>A</u>TG8 to <u>S</u>ingle-<u>M</u>embranes of the endolysosomal system (CASM) (Durgan and Florey, 2022). CASM is implicated in a wide range of processes, including phagocytosis, endocytosis and response to infection (viral, bacterial, yeast), with important functions in immunity (Heckmann et al., 2017) and neurodegeneration (Heckmann et al., 2020; Heckmann et al., 2019). Mechanistically, CASM is independent of canonical autophagy and does not require the ULK/ATG13/FIP200 initiation complex (Florey et al., 2015; Jacquin et al., 2017; Martinez et al., 2015). Instead, CASM engages the ATG16L1 complex to promote ATG8 lipidation directly onto single, endolysosomal membranes, via conjugation to PE, and also to phosphatidylserine (PS); ATG8-PS conjugation is a specific molecular signature of CASM (Durgan et al., 2021). A broad set of stimuli can activate CASM, including ROS generation, pathogens, lysosomotropic drugs, ionophores, activation of the STING pathway and agonists of the lysosomal Ca2+ channel TRPML1. These diverse stimuli all share the common feature of perturbing the ionic and pH balance of the endolysosome system (Durgan and Florey, 2022). In response to elevated lysosomal pH, the V1 and V0 sectors of V-ATPase engage at the lysosome membrane, to drive reacidification. This enhanced engagement also directly recruits ATG16L1, specifying the site for ATG8 lipidation by CASM. This V-ATPase-ATG16L1 (VAIL) axis is dependent on critical residues in the ATG16L1 C-terminal WD40 domain, including K490 (Fletcher et al., 2018), and can be inhibited by the *Salmonella* effector protein, SopF (Fischer et al., 2020; Hooper et al., 2022; Ulferts et al., 2021; Xu et al., 2019). While the functional significance of the CASM pathway is now well established, the precise downstream mechanisms remain to be fully defined. Recent work has defined a CASM pathway involved in lysosome biogenesis, and this field lies open for further investigation.

Interestingly, some recent studies have suggested that an unconventional form of ATG8 lipidation may occur at damaged lysosomes (Jia et al., 2022; Kumar et al., 2020; Nakamura et al., 2020; Tan and Finkel, 2022; Xu et al., 2022). Here, we tested the hypothesis that CASM may provide a mechanistic explanation, undertaking a comprehensive analysis of ATG8 lipidation in the early phases of lysosome damage. Using a panel of complementary biochemical, pharmacological and genetic approaches, we show that the robust ATG8 lipidation observed at lysosomes rapidly after damage indeed occurs via the CASM pathway. Furthermore, we provide evidence that lysosome damage-induced CASM engages the lipid transfer protein ATG2, building further links between this protein and lysosome homeostasis. Finally, we discover that ATG2 engagement represents a shared downstream response to a range of CASM stimuli, suggesting this pathway may a broader role in the response to stress at various membranes.

## Results and Discussion

### Rapid ATG8 lipidation in response to lysosome damage is largely independent of canonical autophagy

To visualise early ATG8 dynamics during lysosome damage, we used live cell imaging to monitor GFP-LC3A in MCF10A cells. Addition of lysosome damaging agents, L-leucyl-leucine methyl ester (LLOMe) or glycyl-l-phenylalanine 2-naphthylamide (GPN), induced rapid and robust relocalisation of GFP-LC3A, in less than 12 minutes (Fig. 1A, Video 1 and 2). Consistent with this, GFP-LC3A can be detected by western blotting upon LLOMe treatment, with the appearance of the characteristic, band-shifted LC3-II form (Fig. 1B). To investigate the mechanism underlying early ATG8 lipidation upon lysosome damage, we used genetic and pharmacological approaches to inhibit of canonical autophagy. ATG13KO cells, which are deficient for canonical autophagy, or wild type (WT) controls, were treated with LLOMe, to promote lysosome damage, or with PP242 as a control to induce canonical autophagy. As expected, only WT cells responded to PP242 (Fig. 1C and D). However, both WT and ATG13 KO cells displayed robust GFP-LC3A relocalisation in response to LLOMe, suggesting this response occurs independently from canonical autophagy. Consistent with this, inhibition of canonical autophagy, using a Vps34 inhibitor (IN-1), or chelation of intracellular calcium (BAPTA-AM), blocked the GFP-LC3A response to PP242, but had no effect on LLOMe induced GFP-LC3A relocalisation (Fig. 1C and D), further indicating independence from canonical autophagy. Finally, as a complementary approach, we assessed the dynamics of an early autophagosome marker, WIPI2. As expected, PP242 stimulation induced WIPI2 puncta in WT cells, but not ATG13 KO. Similarly, LLOMe induced a small increase in WIPI2 puncta in WT cells, but also had no effect in ATG13 KO cells (Fig. 1E and F), further confirming that the LC3A response to LLOMe occurs largely in the absence of any autophagosome formation at this timepoint.

**Figure 1.**
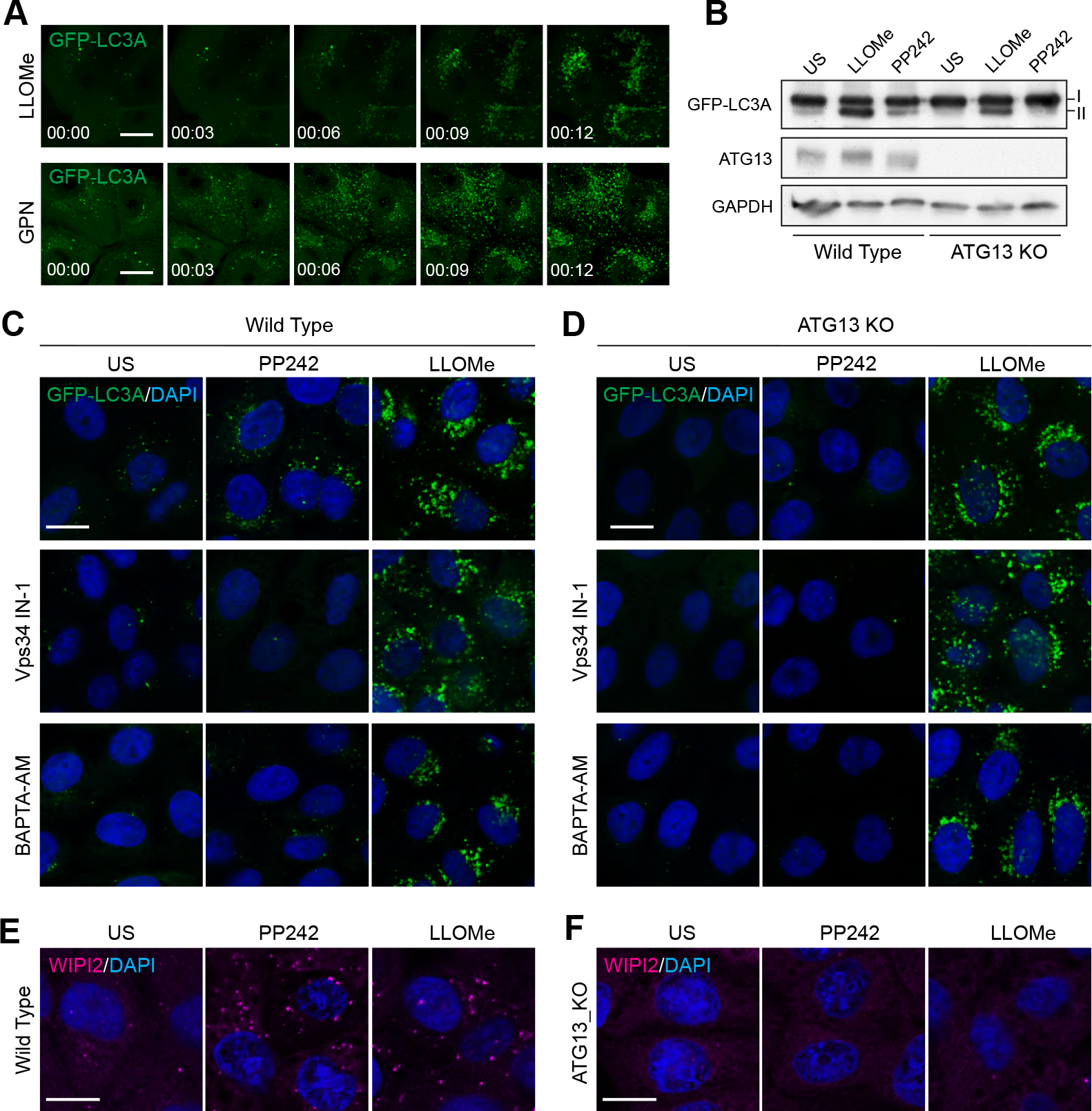
Lysosome damage induces LC3A lipidation independent of canonical autophagy. (A) Representative time-lapse confocal z stack images of MCF10A cells expressing GFP-LC3A treated with 250 µM LLOMe or GPN 200 µM. Scale bar, 10 µm; hr:min. (B) Wild type and ATG13 KO MCF10A cells expressing GFP-LC3A were treated with LLOMe (250 µM, 20 min) or PP242 (1 µM, 1 hr). Western blotting was performed to probe for GFP-LC3A (I and II forms are marked). Representative confocal images of GFP-LC3A in (C) wild type and (D) ATG13 KO MCF10A cells treated with LLOMe (250 µM, 20 min) or PP242 (1 µM, 1 hr), after pre-treatment with Vps34 IN-1 (5 µM) or BAPTA-AM (10 µM). Scale bar, 10 µm. Representative confocal images of (E) wild type and (F) ATG13 KO MCF10A cells treated as in (C) and (D), and stained for WIPI2. Scale bar, 10 µm.

Together these data clearly demonstrate that early ATG8 lipidation induced by lysosome damage is largely independent from canonical autophagy; these findings align with observations from other recent work (Jia et al., 2022; Kumar et al., 2020; Nakamura et al., 2020; Tan and Finkel, 2022; Xu et al., 2022). While conventional lysophagy occurs, this represents only a minor portion of the observed ATG8 lipidation at early timepoints, suggesting that multiple autophagy-related pathways drive temporally and mechanistically distinct ATG8 responses to lysosome damage.

### LC3A is recruited directly to damaged lysosomes and is associated with their tabulation

While GFP-LC3A did not co-localise with WIPI2 during early lysosome damage, we observed strong co-localisation with the lysosome marker, LAMP1, in response to LLOMe (Fig. 2A). The degree of co-localisation under these conditions was even greater than that detected during canonical autophagy induction, where LC3 is expected to localise to both LAMP1-negative autophagosomes and LAMP1-positive autophagolysosomes. These data suggest that LC3A is directly recruited to the lysosome membrane upon LLOMe treatment.

**Figure 2.**
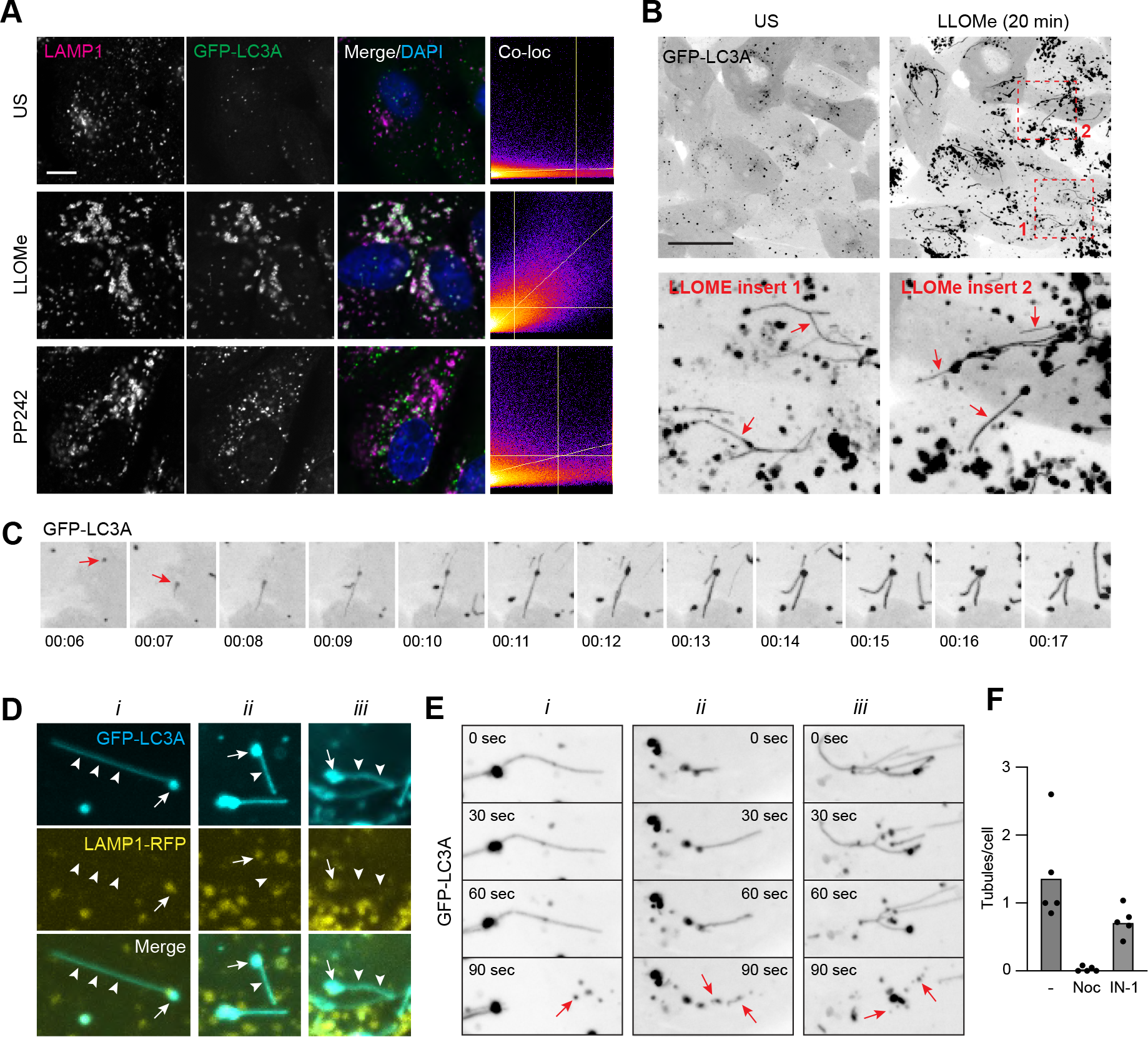
LLOMe induces LC3 recruitment directly to lysosomes and their tubulation. (A) Representative confocal images of MCF10A cells expressing GFP-LC3A, treated with LLOMe (250 µM, 20 min) or PP242 (1 µM, 1 hr) and stained for LAMP1. Co-localization analysis is shown. Scale bar, 5 µm. (B) Images from time-lapse confocal microscopy before and after 20 mins LLOMe (250 µM) treatment. Inserts show 2 areas of LLOMe treated cells. Arrows denote GFP-LC3A positive tubules. Scale bar, 30 µm. (C) Images from time-lapse series showing the formation of GFP-LC3A puncta following LLOMe (250 µM) treatment (red arrow) and subsequent tubulation. min:sec. (D) MCF10A cells co-expressing GFP-LC3A and RFP-LAMP1 treated with LLOMe (250 µM). Images show three examples (*i, ii, iii*). Arrows denote puncta and arrowheads denote tubules. (E) Time-lapse confocal images of LLOMe induced GFP-LC3A tubulation. Three examples are shown (*i, ii, iii*), red arrows denote vesiculation of tubules. (F) Quantification of tubules in cells treated with LLOMe (250 µM). Number of tubules per cell were analysed in 5 fields of view from cells pre-treated with nocodazole (1 µM) or Vps34 IN-1 (5 µM). A total of >150 cells per condition were analysed.

To examine this process further, live cell imaging studies were performed, tracking GFP-LC3A localisation shortly after lysosome damage. These experiments uncovered a striking and highly dynamic pattern of GFP-LC3A localisation (Fig. 2B and C, Video 3 and 4), not preserved in fixed samples. We observed the formation of highly dynamic, GFP-LC3A positive tubules, which appear to emanate from GFP-LC3A puncta. Using a LAMP1-RFP reporter, we found that while the GFP-LC3A-positive puncta are also positive for LAMP1, the distinctive GFP-LC3A tubules are LAMP1 negative (Fig. 2D). Strikingly, the tubules frequently undergo scission events, resulting in the formation of smaller GFP-LC3A positive vesicles (Fig. 2E, Video 5 and 6). Tubule formation and movement is blocked by nocodazole, suggesting a requirement for microtubules (Fig. 2F). The tubules still formed in ATG13 KO cells, and did not increase with Vps34 inhibition, so seem unlikely to be related to Autophagic Lysosome Reformation (ALR) (Munson et al., 2015) (Fig. 2F). This is reminiscent of tubule formation seen during LYTL (Bonet-Ponce et al., 2020), where, however, no involvement of ATG8 lipidation had been reported.

Together, these data indicate that, during the early phases of lysosome damage, ATG8 is lipidated directly onto lysosomes, from where it can be incorporated into highly dynamic tubular structures.

### Lysosome damage induced ATG8 conjugation bears the hallmarks of CASM

We next considered the possible molecular mechanisms underlying the rapid and direct recruitment of ATG8 to lysosomes upon damage. Given the process is independent of canonical autophagy, and involves targeting of single membrane endolysosomes, we reasoned that CASM represented a strong candidate mechanism, and tested this possibility using a series of biochemical, pharmacological and genetic approaches.

Firstly, we assessed the molecular profile of ATG8 lipidation associated with lysosome damage. CASM, unlike canonical autophagy, drives the alternative conjugation of ATG8 to phosphatidylserine (PS), providing a diagnostic molecular feature to distinguish between these related autophagy pathways (Durgan et al., 2021). To characterise ATG8 lipidation following lysosome damage, mass spectrometric analysis of GFP-LC3A was undertaken, in WT cells -/+ LLOMe treatment. Importantly, LLOMe induced the formation of GFP-LC3A-PS, as well as GFP-LC3A-PE (Fig. 3 A and B), consistent with the signature of CASM. V-ATPase is a critical player in CASM, and inhibition of V-ATPase with Bafilomycin A1 (BafA1) characteristically blocks this pathway (Florey et al., 2015; Jacquin et al., 2017), while simultaneously elevating levels of canonical autophagosomes. We found that BafA1 reduced the formation of both GFP-LC3A-PS, and GFP-LC3A-PE, following LLOMe treatment, again consistent with the hallmark features of CASM.

**Figure 3.**
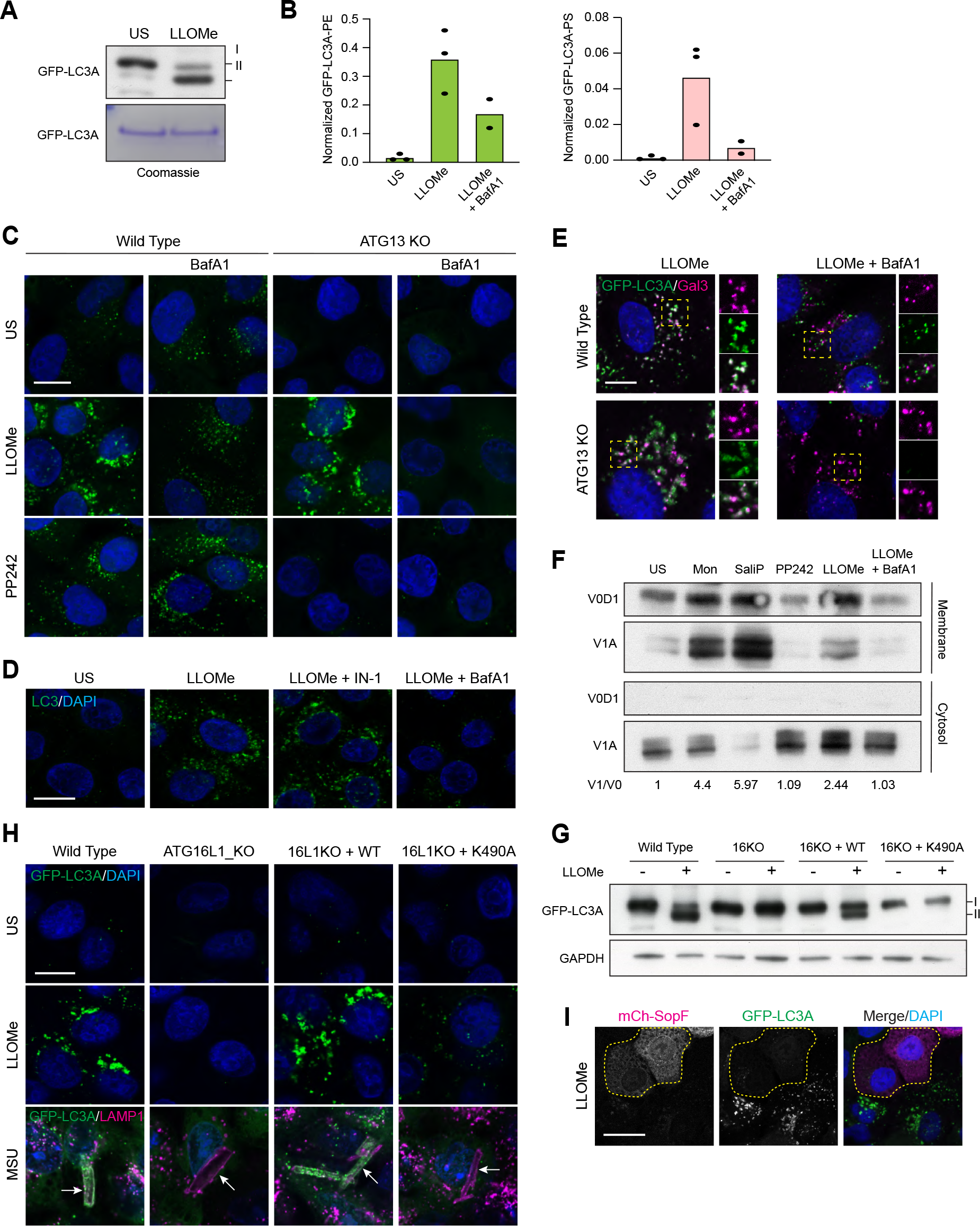
Lysosome damage activates CASM through the V-ATPase-ATG16L2 axis. (A) Western blotting and Coomassie staining of GFP-IPs from MCF10A GFP-LC3A expressing cells treated +/- LLOMe (250 µM, 40 mins). (B) Normalized mass spectrometry analysis of LC3A-PE and LC3A-PS in cells treated as in (A). (C) Representative confocal images of GFP-LC3A in wild type and ATG13 KO MCF10A cells treated with LLOMe (250 µM, 30 min) or PP242 (1 µM, 1 hr), after pre-treatment with BafA1 (100 nM). Scale bar, 10 µm. (D) Representative confocal images of endogenous LC3 in wild type MCF10A cells treated as in (C). Scale bar, 10 µm. (E) Confocal images of wild type or ATG13 KO MCF10A cells expressing GFP-LC3A treated with LLOMe (250 µM, 20 min) +/- BafA1 pre-treatment (100 nM) and stained for Galectin-3. Inserts show single channel cropped images. Scale bar, 5 µm. MCF10A cells were treated with monensin (100 μM), SaliP (2.5 μM), PP242 (1 µM), LLOMe (250 μM) or LLOMe + BafA1 (100 nM) for 1 h. Following fractionation, membrane and cytosol fractions were probed for ATP6V1A, ATP6V0D1 by western blotting. V1/V0 ratios shown below. (G) Western blotting analysis of GFP-LC3A expressing wild type MCF10A and ATG16L1 KO cells re-expressing ATG16L1 WT or K490A, treated with LLOMe (250 µM, 20 min). (H) Confocal images of GFP-LC3A in the MCF10A ATG16L1 cell line panel treated with LLOMe (250 µM, 20 mins) or MSU crystals (200 µg/ml, 4 hr) and co stained for LAMP1. Arrows denote LAMP1 positive crystals. Scale bar, 10 µm. (I) Confocal images of GFP-LC3A expressing MCF10A cells co-transfected with mCherry-SopF. Outlined cells mark mCherry-SopF expressing cells. Scale bar, 20 µm.

To interrogate this mechanism further, we combined pharmacological and genetic approaches, to disrupt CASM or canonical autophagy, for imaging-based analyses. In WT MCF10A cells, BafA1 dramatically reduced the GFP-LC3A response to LLOMe, consistent with CASM, while having the expected, opposing effect of increasing PP242-induced GFP-LC3A puncta, by inhibiting canonical autophagic flux (Fig. 3C). Moreover, in ATG13 KO cells, which lack basal autophagosomes, BafA1 completely blocked the GFP-LC3A response to LLOMe (Fig. 3C). In agreement with these findings, we found that LLOMe similarly induces endogenous LC3 puncta, in a manner blocked by BafA1, but unaffected by Vps34 inhibition (Fig. 3D). Importantly, the inhibitory effects of BafA1 cannot be attributed to suppression of lysosome damage, as galectin 3 puncta still formed (Fig. 3E). Interestingly, in WT cells, BafA1 reduced the colocalization of GFP-LC3A with Galectin 3, suggesting that, in these cells, the residual GFP-LC3A puncta represent background autophagosomes.

In a final, complementary approach to exclude the input of canonical autophagy, we analysed a more precisely defined endolysosomal compartment, completely unrelated to autophagosomes. Using latex bead engulfment to induce phagocytosis, we observed LLOME-induced recruitment of GFP-LC3A to LAMP1-positive, latex bead containing phagosomes (Fig. S1A and B), again in BafA1-sensitive manner.

Together, these data confirm that LLOMe-induced ATG8 lipidation bears several hallmark features of CASM, including the molecular signature of ATG8-PS, and the distinctive pharmacological profile of BafA1 sensitivity, Vps34 insensitivity.

### Lysosome damage harnesses the V-ATPase-ATG16L1 axis

A shared feature of CASM stimuli is disruption of endolysosomal ion and pH balances, which in turn drive increased engagement of V-ATPase V0-V1 sectors (Hooper et al., 2022). We reasoned that if lysosome damage drives ATG8 lipidation via CASM, it would also be expected to increase V0-V1 engagement. Using membrane fractionation and western blotting, translocation of the cytosolic V1A subunit to membranes was measured as a read-out. Strikingly, LLOMe treatment, similar to known inducers of CASM such as monensin and SaliPhe, increased translocation of the cytosolic V1A subunit to membranes, in a BafA1-sensitive manner (Fig. 3F).

The next step in CASM involves the V-ATPase-ATG16L1 axis, through which increased V0-V1 association promotes the direct recruitment of ATG16L1, via its C-terminal WD40 domain, dependent upon the K490 residue (Hooper et al., 2022; Ulferts et al., 2021). An ATG16L1 K490A mutant is deficient for CASM, but competent for canonical autophagy, providing a specific genetic tool with which to dissect these pathways. Using a panel of cells, including WT, ATG16L1 KO, and KOs re-expressing either WT or K490A ATG16L1, we analysed the molecular response to lysosome damage. ATG16L1 KO cells do not undergo GFP-LC3 lipidation upon LLOMe treatment, as detected by western blotting, and this feature can be restored by expression of wild type ATG16L1, but not the CASM deficient K490A mutant (Fig. 3G). Similar K490 dependency was observed in the GFP-LC3A response to LLOMe using immunofluorescence confocal imaging (Fig. 3H). Importantly, we also found that when lysosome damage is induced by Monosodium Urate (MSU) crystals, a more physiological stimulus, recruitment of GFP-LC3A to LAMP1-positive MSU containing compartments was similarly dependent on ATG16L1 K490 (Fig. 3H).

Finally, the interaction between V-ATPase and ATG16L1 during CASM is inhibited by the *Salmonella* effector protein, SopF (Xu et al., 2019), which acts in concert with ARF1 to ribosylate the V0c subunit of V-ATPase. Strikingly, we found that expression of mCherry-SopF substantially impaired LLOMe-induced GFP-LC3A puncta, as judged by fluorescent microscopy (Fig. 3I).

Together, these findings demonstrate comprehensively that lysosome damage, triggered by multiple stimuli, potently activates CASM, via the V-ATPase-ATG16L1 axis, as the primary early ATG8 response.

### Lipidated LC3A interacts with ATG2 during CASM

ATG8 proteins perform a range of important functions in autophagy-related processes, often dependent on interaction with proteins that harbour LC3 interacting regions (LIR). In canonical autophagy, LIR-based binding is critical for both cargo recognition and maturation of autophagosomes (Johansen and Lamark, 2020). Surprisingly little is known about the downstream molecular mechanisms of CASM, and a key open question is whether ATG8 proteins similarly recruit effector proteins to endolysosomal membranes.

ATG2 isoforms A and B, are lipid transfer proteins which undergo LIR-dependent interaction with ATG8s, specifically GABARAPs and LC3A isoforms (Bozic et al., 2020). They play a critical role in supplying lipids to support autophagosome expansion (Osawa et al., 2019; Valverde et al., 2019) during canonical autophagy, and have recently been implicated in lysosome damage repair through the PITT pathway (Tan and Finkel, 2022). In light of these observations, and the data presented so far, we questioned whether CASM may involve ATG2 engagement during lysosome damage. Using GFP-TRAP immunoprecipitation, we found that LLOMe indeed induced a robust interaction between GFP-LC3A and ATG2B (Fig. 4A). Importantly, this interaction is independent of canonical autophagy, as it still occurred in ATG13 KO cells (Fig. 4A). Furthermore, while Vps34 inhibition blocked the GFP-LC3A ATG2B interaction induced by PP242 (canonical autophagy), it did not block the LLOMe-induced interaction (Fig. 4B). The interaction with ATG2B is dependent on GFP-LC3A lipidation, as it was lost in ATG16L1 KO cells, and more specifically on CASM, because it could be rescued by WT ATG16L1, but not K490A expressing cells (Fig. 4C). Thus, lysosome damage appears to promote the engagement of ATG2, via the CASM specific interaction with ATG8 proteins on the lysosome membrane. Consistent with this model, we observed increased amounts of ATG2B in membrane fractions with LLOMe treatment, which was dependent on ATG16L1, but not ATG13 (Fig. 4D).

**Figure 4.**
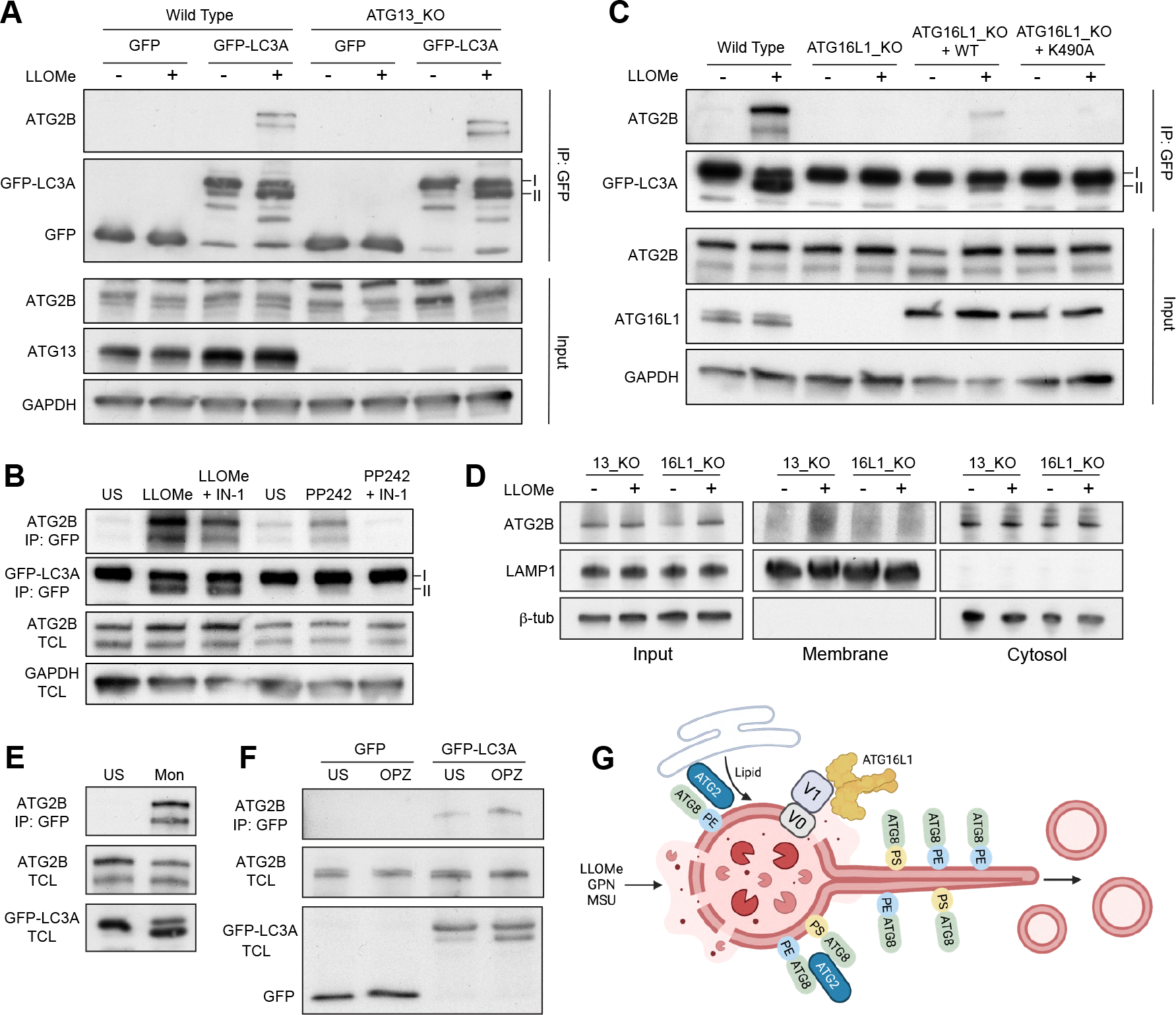
LC3A interacts with ATG2B during lysosome-induced CASM. (A) Western blot analysis of GFP-IPs and input from wild type and ATG13 KO MCF10A cells expressing either GFP or GFP-LC3A treated +/- LLOMe (250 µM, 30 mins). Samples probed for ATG2B, GFP, ATG13 and GAPDH. (B) Western blot analysis of GFP-IPs and input from MCF10A ATG16L1 cell line panel expressing GFP-LC3A treated +/- LLOMe (250 µM, 30 mins). Samples probed for ATG2B, GFP, ATG16L1 and GAPDH. (C) Western blot analysis of GFP-IPs and total cell lysate (TCL) from GFP-LC3A expressing MCF10A cells treated with LLOMe (250 µM, 30 mins) or PP242 (1 µM, 1 hr) +/- Vps34 IN-1 (5 µM). Samples probed for ATG2B, GFP and GAPDH. (D) GFP-LC3A expressing ATG13 or ATG16L1 KO MCF10A cells were treated with LLOMe (250 µM, 30 mins). Following fractionation, input, cytosol and membrane fractions were probed for ATG2B, LAMP1, GFP and β-tubulin. (E) Western blot analysis of GFP-IPs and input from ATG13 KO MCF10A cells expressing GFP-LC3A treated with LLOMe (250 µM, 30mins), GPN (200 µM, 30 min) or monensin (100 µM, 40 mins). Samples probed for ATG2B and GFP. (F) Western blot analysis of GFP-IPs and input from RAW264.7 cells expressing GFP-LC3A following incubation with opsonized zymosan (OPZ) for 40 mins. Samples were probed for ATG2B and GFP. (G) Model of ATG8 response to lysosome damage illustrating the ATG8 associated lysosome tubulation and vesiculation, activation of the V-ATPase-ATG16L1 axis, ATG8 conjugation to PE and PS, and the engagement of lipid transfer protein ATG2. Image created using Biorender.

Together, these data indicate that CASM drives an ATG8-ATG2 interaction, in response to lysosome damage. This finding provides important new molecular insights into both the downstream consequences of CASM, and the central role of ATG2 in lysosome homeostasis.

In considering these findings, we wondered whether the inducible engagement of the ATG2 lipid transporter may represent a broader feature of CASM, which has been conceptualised as a response to membrane stress (Kumar et al., 2021). To test this, we assessed the ability of other CASM stimuli to induce a GFP-LC3A ATG2B interaction. Strikingly, other CASM stimuli, including monensin (lysosomotropic) and LC3-associated phagocytosis (LAP) in RAW264.7 cells all induced a GFP-LC3A-ATG2B interaction (Fig. 4E and F). Together, these findings suggest that CASM drives ATG2 engagement as a general response to lysosomal perturbation.

Through this work, we have established clearly that CASM is activated in rapid response to perturbation of lysosomal permeability (Fig. 4G). We suggest that CASM underlies the ‘unconventional’ ATG8 lipidation observed following lysosome damage in some recent studies. In parallel, classical lysophagy acts to clear away the damaged organelles, suggesting that multiple, distinct autophagy-related pathways contribute to maintenance of lysosome integrity. Considering that CASM and lysophagy share the common ATG8 conjugation machinery, caution is required when interpreting results from autophagy protein knockout or ATG8 lipidation experiments in this context (and others). Our current data establish comprehensively that ATG8 proteins can be conjugated directly onto lysosome membranes by CASM, and can recruit an effector protein, ATG2, in an analogous manner to canonical autophagy. However, this interacting protein will not be cargo destined for degradation, but may instead exert adapter function directly at the lysosome membrane. Notably, another ATG8 protein, GABARAP, can engage with FNIP/Folliculin at lysosome membranes, during TRPML1-agonist induced CASM, acting as an adapter to promote TFEB-mediated lysosome biogenesis (Goodwin et al., 2021). We now reveal that LC3A engages with ATG2B at damaged lysosomes. ATG2 plays an important role in lysosome repair through the recently described PITT pathway (Tan and Finkel, 2022). Although PITT occurs independently of ATG8 lipidation and autophagy, we have now identified a CASM-dependent engagement of ATG2, that may run parallel to PITT, stabilising or reinforcing its function. As fundamental cellular processes often engage multiple, parallel pathways to safeguard proper function and integrate complex signalling information, we propose that CASM may represent one of several, co-operative mechanisms responding to lysosome damage, involving ATG8s and/or ATG2s. Furthermore, because other CASM stimuli similarly induce ATG2 engagement, we speculate that CASM may act as part of a more generalised mechanism to maintain endolysosomal homeostasis in response to stress, by engaging with lipid transport machinery.

## Acknowledgements

We would like to thank all members of the Florey Lab for their help. We would also like to thank the Imaging and Mass Spectrometry facilities at the Babraham Institute for their support and input. This work was supported by grants from the BBSRC, BB/P013384/1 (BBS/E/B/000C0432 and BBS/E/B/000C0434), and a Cambridge MRC DTP studentship. O. Florey is an ad hoc paid consultant for Casma Therapeutics. The authors declare no further competing financial interests

## Contributions

J.C. designed and carried out and analysed experiments. D.G.M. provided key reagents and advice. J.D. designed experiments and wrote the paper. O.F. designed, carried out and analysed experiments and wrote the manuscript.

## Methods

### Antibodies

Antibodies used were rabbit polyclonal anti-ATG16L1 (8090, Cell Signalling, WB 1:1000), rabbit polyclonal anti-LC3A/B (4108, Cell Signalling, WB 1:1000, IF 1:100), rabbit monoclonal anti-ATP6V1A (ab199326, Abcam, WB 1:2000), mouse monoclonal anti-ATP6V0d1 (ab56441, Abcam, WB 1:1000), mouse monoclonal anti-LAMP1 (555798, BD Biosciences, IF 1:100), mouse monoclonal anti LAMP1 (611042, BD Bioscience, WB, 1:500), mouse monoclonal anti-GAPDH (ab8245, Abcam, WB 1:1000), rabbit anti-ATG13 (6940, Cell Signalling, WB 1:1000), mouse anti-GFP (1181446000, Roche, WB 1:1000), mouse anti-WIPI2 (MCA5780GA, BioRad, IF 1:100), rabbit anti ATG2B (25155-1-AP, ProteinTech, WB 1:1000), mouse anti-Galectin 3 (32790, Scbt, IF 1:100), rabbit anti-b tubulin (2128, Cell Signalling, WB 1:1000), Alexa Fluor 488 polyclonal goat anti-rabbit IgG (A-11034, ThermoFisher, IF 1:500), Alexa Fluor 568 polyclonal goat anti-mouse IgG (A-11004, ThermoFisher, IF 1:500), Alexa Fluor 568 polyclonal goat anti-rabbit IgG (A-11011, ThermoFisher, IF 1:500), HRP-conjugated anti-rabbit IgG (7074, Cell Signalling, WB 1:2000), HRP-conjugated anti-mouse IgG (7076, Cell Signalling, WB 1:2000).

### Reagents

Reagents and chemicals used were LLOMe (L7393, Sigma,), GPN (SC-252858, Scbt), BafA1 (1334, Tocris), PP242 (4257, Tocris), Monensin (M5273, Sigma), human serum (H2918, Sigma), Zymosan (Z4250, Sigma), murine IFNγ (315-05, Peprotech), DAPI (D9542, Sigma), IN-1 (17392, Caymen), Saliphenyhalamide (SaliP) and TRPML1 agonist C8 were kindly provided by Casma Therapeutics, USA. BAPTA-AM (B1205, Thermo Fisher), nocodazole (M1404, Sigma), GFP-Trap (gtma-20) and control magnetic agarose beads (bmab-20) were obtained from Chromotek,

### Cell Culture

WT, ATG13 KO and ATG16L1 KO MCF10A cells (human breast epithelial), expressing GFP-LC3A (human), were prepared as described previously (Fletcher et al., 2018; Jacquin et al., 2017) and cultured in DMEM/F12 (Gibco, 11320074) containing 5% horse serum (ThermoFisher, 16050-122), EGF (20 ng/ml; Peprotech AF-100-15), hydrocortisone (0.5 mg/ml; Sigma, H0888), cholera toxin (100 ng/ml; Sigma, C8052), insulin (10 µg/ml; Sigma, I9278), and penicillin/streptomycin (100 U/ml; Gibco 15140-122) at 37°C, 5% CO_2_.

RAW264.7 cells were obtained from ATCC and maintained in DMEM (Gibco, 41966-029) supplemented with 10% FBS (Sigma, F9665) and penicillin/streptomycin (100 U/ml, 100 µg/ml; Gibco 15140-122) at 37°C, 5% CO_2_.

### Retrovirus generation and infection

For Flag-S-ATG16L1 virus infection, MCF10A cells were seeded in a 6 well plate at 5 × 10^4^ per well. The next day 1 ml viral supernatant was added with 10 µg/ml polybrene for 24 h followed by a media change. Cells were then selected with puromycin (2µg/ml).

### Fixed immunofluorescent confocal microscopy and analysis

Cells were seeded in 12-well plates containing coverslips and incubated at 37°C, 5% CO_2_ for 24 h. Following treatments, cells were washed twice with ice cold PBS then incubated with 100% methanol at -20°C for 10 min. The cells were then washed twice with PBS and blocked with 5% BSA (Sigma, A7906) in PBS for 1 h at room temperature. The cells were incubated overnight at 4°C with the primary antibodies, then washed x3 in cold PBS. Fluorescent secondary antibodies were used at a 1:500 dilution in PBS + 5% BSA and were incubated with the cells for 1 h at room temperature. The cells were washed x3 in cold PBS prior to being incubated with DAPI for 10 min at room temperature and then mounted onto microscope slides with ProLong Gold anti-fade reagent (Invitrogen, P36930). Image acquisition was made using a Zeiss LSM 780 confocal microscope (Carl Zeiss Ltd), using Zen software (Carl Zeiss Ltd).

### Live cell confocal microscopy

Cells were seeded into 35 mm glass bottom dishes (MatTek). Images stacks were acquired every 30 seconds using an Olympus SpinSR confocal microscope, comprising Olympus IX83 stand, Olympus 60x 1.5 NA UPLAPO objective lens, Yokogawa CSU-W1 scanhead and Hamamatsu Orca Fusion camera. Images were acquired with a 2×2 camera bin, giving a pixel size of 108 nm. Laser excitation and emission filters for the GFP and RFPchannels were 488 nm (ex) 525/50 nm (em) and 561 nm (ex) 617/73 nm (em) respectively. The image acquisition software was Olympus cellSens v4.1.

### Cell Lysis and GFP-TRAP Immunoprecipitation

Cells expressing GFP-LC3A were seeded across multiple 15-cm dishes, treated as indicated, then placed on ice and washed with ice-cold PBS. Each 15-cm dish was scraped into 900 μl lysis buffer (50 mM Tris pH 7.5, 150 mM NaCl, 0.5% NP40 (IGEPAL CA-630, Sigma I3021), phosphatase inhibitors (1x, Sigma P0044) and protease inhibitors (1x, Sigma P8340). The resulting suspension was incubated on ice for 10 minutes, then centrifuged at 16000 rcm, 4°C, 10 minutes to separate the pellet from the soluble lysate. A small fraction of the supernatant was removed for western blotting, as described below, and the remaining lysate subjected to preclearing and IP, using magnetic beads (Chromotek) and a magnetic separation rack (Cell Signalling), according to the manufacturers’ instructions. The lysate was pre-cleared, using 10 μl equilibrated magnetic agarose control beads/sample (bmab, Chromotek), for 30 minutes, 4°C, on a rotating wheel. Cleared lysates were then incubated with 10 μl equilibrated GFP-TRAP beads/sample (gtma, Chromotek) for 60 minutes, 4°C, on a rotating wheel, to recover GFP-LC3A. The beads were washed 3 × 10 minutes in lysis buffer at 4°C, on a rotating wheel. Enriched GFP-LC3A was eluted for analysis by western blotting or mass spectrometry with the addition of 25 μl 2x LDS buffer (Invitrogen)/0.2 M DTT sample buffer at 100°C, 5 minutes.

For other samples, cells were scraped into ice-cold RIPA buffer (150 mM NaCl, 50 mM Tris–HCl, pH 7.4, 1 mM EDTA, 1% Triton X-100 (Sigma, T8787), 0.1% SDS (Sigma, L3771), 0.5% sodium deoxycholate (Sigma, D6750) and lysed on ice for 10 min. Lysates were centrifuged for 10 min at 10,000 *g* at 4°C.

### Membrane Fractionation

MCF10A cells were seeded on a 15 cm dish and cultured for 48 h. Cells were stimulated in suspension as indicated for 40 mins. Input, cytosol and membrane fractions were isolated using the Mem-Per Plus Membrane Protein Extraction Kit (89842, Thermofisher) following product guidelines. Protein concentration was measured by BCA assay and equal amounts loaded onto polyacrylamide gels for SDS–PAGE analysis.

### Mass Spectrometric Analysis of Lipidated ATG8

GFP-LC3A samples were analysed by mass spectrometry as described in (Durgan et al., 2021). In brief, samples were run on 10% NuPAGE gels in MOPS buffer (Invitrogen). Each gel was washed stained with Imperial Stain (Thermo Scientific, 24615) for 2 hrs, then destained in dH2O overnight. The gel region containing GFP-LC3A was excised into a single tube, destained, and typically saponified by treatment with 50 mM NaOH in 30% MeOH at 40°C for 2 hr. The protein was digested with AspN protease (Roche) at 30°C for 16 hr, in 25 mM ammonium bicarbonate. For targeted mass spectrometric assay of modified C-terminal LC3A peptides, samples were separated on a reversed-phase nanoLC column (150 × 0.075mm; Reprosil-Pur C18AQ, Dr Maisch), interfaced to a Q-Exactive mass spectrometer (Thermo Scientific) operating in high resolution (orbitrap) MS1 mode, with data-dependent acquisition of low resolution MS2 spectra generated by CID in the linear ion-trap.

Quantitative data were extracted using Skyline software (MacCoss Lab, University of Washington) using the sum of the chromatographic peak areas from the y1 to y10 fragment ions. Subsequently, normalization was performed against unmodified C-terminal peptide.

### Western Blotting

Western blotting was performed as described previously (Hooper et al., 2022). Briefly, cell lysates were run on SDS-PAGE gels, transferred to PVDF membrane (Immobilon-P, Millipore), blocked with 5% BSA (Sigma A7906)/TBS-T for 1 hour, RT and then incubated with primary antibody at 4°C overnight. Membranes were washed 3x 10 minutes in TBS-T and incubated with HRP-conjugated secondary antibodies (Cell Signalling 7074, 7076) for 45 minutes, RT. Membranes were washed again 3x 10 minutes in TBS-T, then developed with ECL (GE, RPN2209). Blots were scanned (Epson Perfection, V550).

### Statistics

Unpaired student t-tests, were performed using Graph Pad, as indicated.

## References

Ballabio, A., and J.S. Bonifacino. 2020. Lysosomes as dynamic regulators of cell and organismal homeostasis. Nat Rev Mol Cell Biol. 21:101–118.

Bonet-Ponce, L., A. Beilina, C.D. Williamson, E. Lindberg, J.H. Kluss, S. Saez-Atienzar, N. Landeck, R. Kumaran, A. Mamais, C.K.E. Bleck, Y. Li, and M.R. Cookson. 2020. LRRK2 mediates tubulation and vesicle sorting from lysosomes. Sci Adv. 6.

Bozic, M., L. van den Bekerom, B.A. Milne, N. Goodman, L. Roberston, A.R. Prescott, T.J. Macartney, N. Dawe, and D.G. McEwan. 2020. A conserved ATG2-GABARAP family interaction is critical for phagophore formation. EMBO Rep. 21:e48412.

Durgan, J., and O. Florey. 2022. Many roads lead to CASM: Diverse stimuli of noncanonical autophagy share a unifying molecular mechanism. Sci Adv. 8:eabo1274.

Durgan, J., A.H. Lystad, K. Sloan, S.R. Carlsson, M.I. Wilson, E. Marcassa, R. Ulferts, J. Webster, A.F. Lopez-Clavijo, M.J. Wakelam, R. Beale, A. Simonsen, D. Oxley, and O. Florey. 2021. Non-canonical autophagy drives alternative ATG8 conjugation to phosphatidylserine. Mol Cell. 81:2031–2040.e2038.

Eguchi, T., T. Kuwahara, M. Sakurai, T. Komori, T. Fujimoto, G. Ito, S.I. Yoshimura, A. Harada, M. Fukuda, M. Koike, and T. Iwatsubo. 2018. LRRK2 and its substrate Rab GTPases are sequentially targeted onto stressed lysosomes and maintain their homeostasis. Proc Natl Acad Sci U S A. 115:E9115–e9124.

Fischer, T.D., C. Wang, B.S. Padman, M. Lazarou, and R.J. Youle. 2020. STING induces LC3B lipidation onto single-membrane vesicles via the V-ATPase and ATG16L1-WD40 domain. J Cell Biol. 219.

Fletcher, K., R. Ulferts, E. Jacquin, T. Veith, N. Gammoh, J.M. Arasteh, U. Mayer, S.R. Carding, T. Wileman, R. Beale, and O. Florey. 2018. The WD40 domain of ATG16L1 is required for its non-canonical role in lipidation of LC3 at single membranes. Embo j. 37.

Florey, O., N. Gammoh, S.E. Kim, X. Jiang, and M. Overholtzer. 2015. V-ATPase and osmotic imbalances activate endolysosomal LC3 lipidation. Autophagy. 11:88–99.

Fujita, N., E. Morita, T. Itoh, A. Tanaka, M. Nakaoka, Y. Osada, T. Umemoto, T. Saitoh, H. Nakatogawa, S. Kobayashi, T. Haraguchi, J.L. Guan, K. Iwai, F. Tokunaga, K. Saito, K. Ishibashi, S. Akira, M. Fukuda, T. Noda, and T. Yoshimori. 2013. Recruitment of the autophagic machinery to endosomes during infection is mediated by ubiquitin. J Cell Biol. 203:115–128.

Goodwin, J.M., W.G.t. Walkup, K. Hooper, T. Li, C. Kishi-Itakura, A. Ng, T. Lehmberg, A. Jha, S. Kommineni, K. Fletcher, J. Garcia-Fortanet, Y. Fan, Q. Tang, M. Wei, A. Agrawal, S.R. Budhe, S.R. Rouduri, D. Baird, J. Saunders, J. Kiselar, M.R. Chance, A. Ballabio, B.A. Appleton, J.H. Brumell, O. Florey, and L.O. Murphy. 2021. GABARAP sequesters the FLCN-FNIP tumor suppressor complex to couple autophagy with lysosomal biogenesis. Sci Adv. 7:eabj2485.

Heckmann, B.L., E. Boada-Romero, L.D. Cunha, J. Magne, and D.R. Green. 2017. LC3-Associated Phagocytosis and Inflammation. J Mol Biol. 429:3561–3576.

Heckmann, B.L., B.J.W. Teubner, E. Boada-Romero, B. Tummers, C. Guy, P. Fitzgerald, U. Mayer, S. Carding, S.S. Zakharenko, T. Wileman, and D.R. Green. 2020. Noncanonical function of an autophagy protein prevents spontaneous Alzheimer’s disease. Sci Adv. 6:eabb9036.

Heckmann, B.L., B.J.W. Teubner, B. Tummers, E. Boada-Romero, L. Harris, M. Yang, C.S. Guy, S.S. Zakharenko, and D.R. Green. 2019. LC3-Associated Endocytosis Facilitates β-Amyloid Clearance and Mitigates Neurodegeneration in Murine Alzheimer’s Disease. Cell. 178:536–551.e514.

Herbst, S., P. Campbell, J. Harvey, E.M. Bernard, V. Papayannopoulos, N.W. Wood, H.R. Morris, and M.G. Gutierrez. 2020. LRRK2 activation controls the repair of damaged endomembranes in macrophages. Embo j. 39:e104494.

Hooper, K.M., E. Jacquin, T. Li, J.M. Goodwin, J.H. Brumell, J. Durgan, and O. Florey. 2022. V-ATPase is a universal regulator of LC3-associated phagocytosis and noncanonical autophagy. J Cell Biol. 221.

Ichimura, Y., T. Kirisako, T. Takao, Y. Satomi, Y. Shimonishi, N. Ishihara, N. Mizushima, I. Tanida, E. Kominami, M. Ohsumi, T. Noda, and Y. Ohsumi. 2000. A ubiquitin-like system mediates protein lipidation. Nature. 408:488–492.

Jacquin, E., S. Leclerc-Mercier, C. Judon, E. Blanchard, S. Fraitag, and O. Florey. 2017. Pharmacological modulators of autophagy activate a parallel noncanonical pathway driving unconventional LC3 lipidation. Autophagy. 13:854–867.

Jia, J., F. Wang, Z. Bhujabal, R. Peters, M. Mudd, T. Duque, L. Allers, R. Javed, M. Salemi, C. Behrends, B. Phinney, T. Johansen, and V. Deretic. 2022. Stress granules and mTOR are regulated by membrane atg8ylation during lysosomal damage. J Cell Biol. 221.

Johansen, T., and T. Lamark. 2020. Selective Autophagy: ATG8 Family Proteins, LIR Motifs and Cargo Receptors. J Mol Biol. 432:80–103.

Kumar, S., A. Jain, S.W. Choi, G.P.D. da Silva, L. Allers, M.H. Mudd, R.S. Peters, J.H. Anonsen, T.E. Rusten, M. Lazarou, and V. Deretic. 2020. Mammalian Atg8 proteins and the autophagy factor IRGM control mTOR and TFEB at a regulatory node critical for responses to pathogens. Nat Cell Biol. 22:973–985.

Kumar, S., J. Jia, and V. Deretic. 2021. Atg8ylation as a general membrane stress and remodeling response. Cell Stress. 5:128–142.

Maejima, I., A. Takahashi, H. Omori, T. Kimura, Y. Takabatake, T. Saitoh, A. Yamamoto, M. Hamasaki, T. Noda, Y. Isaka, and T. Yoshimori. 2013. Autophagy sequesters damaged lysosomes to control lysosomal biogenesis and kidney injury. Embo j. 32:2336–2347.

Martinez, J., R.K. Malireddi, Q. Lu, L.D. Cunha, S. Pelletier, S. Gingras, R. Orchard, J.L. Guan, H. Tan, J. Peng, T.D. Kanneganti, H.W. Virgin, and D.R. Green. 2015. Molecular characterization of LC3-associated phagocytosis reveals distinct roles for Rubicon, NOX2 and autophagy proteins. Nat Cell Biol. 17:893–906.

Munson, M.J., G.F. Allen, R. Toth, D.G. Campbell, J.M. Lucocq, and I.G. Ganley. 2015. mTOR activates the VPS34-UVRAG complex to regulate autolysosomal tubulation and cell survival. Embo j. 34:2272–2290.

Nakamura, S., S. Shigeyama, S. Minami, T. Shima, S. Akayama, T. Matsuda, A. Esposito, G. Napolitano, A. Kuma, T. Namba-Hamano, J. Nakamura, K. Yamamoto, M. Sasai, A. Tokumura, M. Miyamoto, Y. Oe, T. Fujita, S. Terawaki, A. Takahashi, M. Hamasaki, M. Yamamoto, Y. Okada, M. Komatsu, T. Nagai, Y. Takabatake, H. Xu, Y. Isaka, A. Ballabio, and T. Yoshimori. 2020. LC3 lipidation is essential for TFEB activation during the lysosomal damage response to kidney injury. Nat Cell Biol. 22:1252–1263.

Osawa, T., T. Kotani, T. Kawaoka, E. Hirata, K. Suzuki, H. Nakatogawa, Y. Ohsumi, and N.N. Noda. 2019. Atg2 mediates direct lipid transfer between membranes for autophagosome formation. Nat Struct Mol Biol. 26:281–288.

Perera, R.M., and R. Zoncu. 2016. The Lysosome as a Regulatory Hub. Annu Rev Cell Dev Biol. 32:223–253.

Radulovic, M., K.O. Schink, E.M. Wenzel, V. Nähse, A. Bongiovanni, F. Lafont, and H. Stenmark. 2018. ESCRT-mediated lysosome repair precedes lysophagy and promotes cell survival. Embo j. 37.

Radulovic, M., E.M. Wenzel, S. Gilani, L.K. Holland, A.H. Lystad, S. Phuyal, V.M. Olkkonen, A. Brech, M. Jäättelä, K. Maeda, C. Raiborg, and H. Stenmark. 2022. Cholesterol transfer via endoplasmic reticulum contacts mediates lysosome damage repair. Embo j. 41:e112677.

Scheffer, L.L., S.C. Sreetama, N. Sharma, S. Medikayala, K.J. Brown, A. Defour, and J.K. Jaiswal. 2014. Mechanism of Ca²+-triggered ESCRT assembly and regulation of cell membrane repair. Nat Commun. 5:5646.

Skowyra, M.L., P.H. Schlesinger, T.V. Naismith, and P.I. Hanson. 2018. Triggered recruitment of ESCRT machinery promotes endolysosomal repair. Science. 360.

Tan, J.X., and T. Finkel. 2022. A phosphoinositide signalling pathway mediates rapid lysosomal repair. Nature. 609:815–821.

Ulferts, R., E. Marcassa, L. Timimi, L.C. Lee, A. Daley, B. Montaner, S.D. Turner, O. Florey, J.K. Baillie, and R. Beale. 2021. Subtractive CRISPR screen identifies the ATG16L1/vacuolar ATPase axis as required for non-canonical LC3 lipidation. Cell Rep. 37:109899.

Valverde, D.P., S. Yu, V. Boggavarapu, N. Kumar, J.A. Lees, T. Walz, K.M. Reinisch, and T.J. Melia. 2019. ATG2 transports lipids to promote autophagosome biogenesis. J Cell Biol. 218:1787–1798.

Wang, F., R. Gómez-Sintes, and P. Boya. 2018. Lysosomal membrane permeabilization and cell death. Traffic. 19:918–931.

Xu, Y., S. Cheng, H. Zeng, P. Zhou, Y. Ma, L. Li, X. Liu, F. Shao, and J. Ding. 2022. ARF GTPases activate Salmonella effector SopF to ADP-ribosylate host V-ATPase and inhibit endomembrane damage-induced autophagy. Nat Struct Mol Biol. 29:67–77.

Xu, Y., P. Zhou, S. Cheng, Q. Lu, K. Nowak, A.K. Hopp, L. Li, X. Shi, Z. Zhou, W. Gao, D. Li, H. He, X. Liu, J. Ding, M.O. Hottiger, and F. Shao. 2019. A Bacterial Effector Reveals the V-ATPase-ATG16L1 Axis that Initiates Xenophagy. Cell. 178:552–566.e520.

Zhen, Y., M. Radulovic, M. Vietri, and H. Stenmark. 2021. Sealing holes in cellular membranes. Embo j. 40:e106922.

