## Supplemental Material for "Lysosome damage triggers direct ATG8 conjugation and ATG2 engagement via CASM"

**Figure S1**

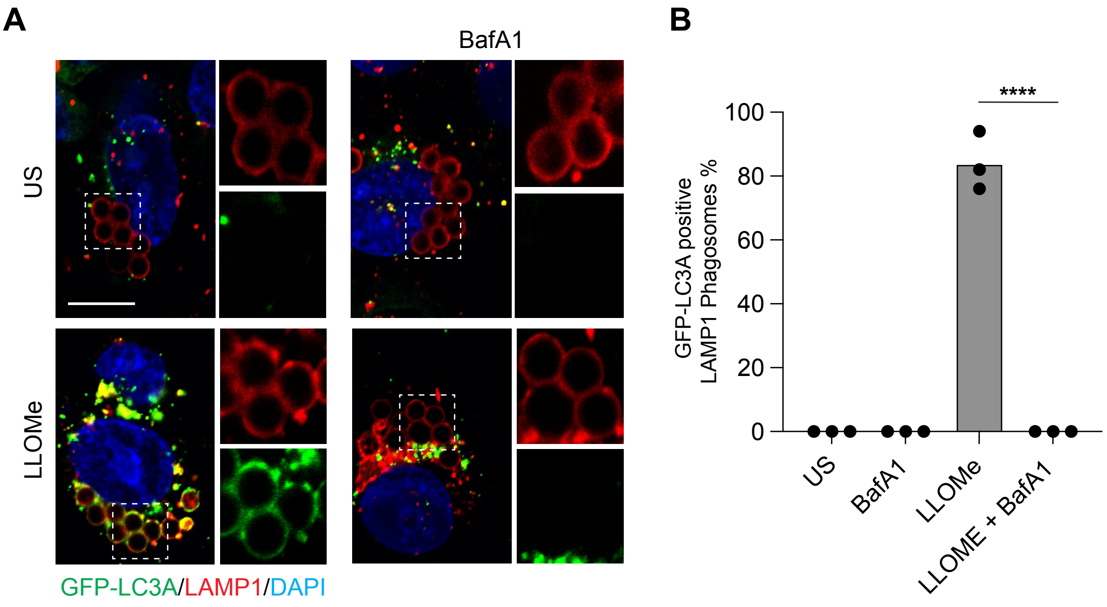


**Figure S1.**

(A) Representative timelapse widefield microscopy images MCF10A cells expressing GFP-LC3A and stained for LAMP1 (red), following engulfment of 3 µm latex beads and treatment with LLOMe (250 µM, 30 mins) +/- BafA1 (100 nM). Cropped images show LAMP1 positive latex- bead containing phagosomes. Scale bar, 10 µm. (B) Quantification of GFP-LC3A recruitment to LAMP1 positive phagosomes. Data represent mean +/- SEM from three independent experiments. ****$p<0.0001$, unpaired t-test.

**Video 1**

Timelapse spinning-disc confocal microscopy showing MCF10A cells expressing GFP-LC3A treated with LLOMe (250 μM). Image stacks acquired every 30 seconds. Movie plays 4 frames/sec; time min:sec. Related to Figure 1A.

**Video 2**

Timelapse spinning-disc confocal microscopy showing MCF10A cells expressing GFP-LC3A treated with GPN (200 μM). Image stacks acquired every 30 seconds. Movie plays 4 frames/sec; time min:sec. Related to Figure 1A.

**Video 3**

Timelapse spinning-disc confocal microscopy showing MCF10A cells expressing GFP-LC3A treated with LLOMe (250 μM). Image stacks acquired every 30 seconds. Movie plays 4 frames/sec; time min:sec. Related to Figure 2B.

**Video 4**

Cropped image sequence from MCF10A cells expressing GFP-LC3A treated with LLOMe (250 μM), showing GFP-LC3A puncta formation and tubulation. Image stacks acquired every 30 seconds. Movie plays 4 frames/sec; time min:sec. Related to Figure 2C.

**Video 5**

Cropped image sequence from MCF10A cells expressing GFP-LC3A treated with LLOMe (250 μM), showing vesiculation of GFP-LC3A tubules. Image stacks acquired every 30 seconds. Movie plays 2 frames/sec; time min:sec. Related to Figure 2E example *iii*.

**Video 6**

Cropped image sequence from MCF10A cells expressing GFP-LC3A treated with LLOMe (250 μM), showing vesiculation of GFP-LC3A tubules. Image stacks acquired every 30 seconds. Movie plays 2 frames/sec; time min:sec. Related to Figure 2E example *ii*.
